# D-Retro Inverso (DRI) Amylin and the Stability of Amylin Fibrils

**DOI:** 10.1101/2020.03.09.984336

**Authors:** Preeti Pandey, Natalie Nguyen, Ulrich H.E. Hansmann

## Abstract

Motivated by the role that amylin aggregates pay in type-II diabetes, we compare the stability of regular amylin fibrils with the stability of fibrils where L-amino acid chains are replaced by D-Retro Inverso (DRI) amylin, *i.e.*, peptides where the sequence of amino acids is reversed, and at the same time the L-amino acids are replaced by their mirror images. Our molecular dynamics simulations show that despite leading to only marginal difference in fibril structure and stability, aggregating DRI-amylin peptides have different pattern of contacts and hydrogen bonding. Because of these differences does DRI-amylin, when interacting with regular (L) amylin, alter the elongation process and lowers the stability of hybrid amylin fibrils. Our results suggest not only a potential use of DRI-amylin as inhibitor of amylin fibril-formation, but points also to the possibility of using insertion of DRI-proteins in L-assemblies as a way to probe the role of certain kinds of hydrogen bonds in supra-molecular assemblies or aggregates.

## 1 Introduction

While presence of amyloid fibrils is most commonly associated with neurodegenerative diseases such as Alzheimer’s, Parkinson’s,or Huntington’s disease, they are also seen in metabolic and other illnesses. One example is diabetes mellitus type-II, where aggregation of islet amyloid polypeptide (IAPP, also known as amylin) in the pancreas causes endoplasmic reticulum stress, mitochondrial damage and membrane disruption.^1,2^ The resulting death of the pancreatic islet *β*-cells, which in healthy persons modulate the secretion of insulin and glucagon, leads to the onset of type-II diabetes.^3–10^ Co-localization of amylin and *β*-amyloid (A*β*) fibrils and computational studies indicate cross-seeding between the two molecules. Hence, inhibition and degradation of amylin aggregates might be a therapeutic strategy not only against type-II diabetes but also against Alzheimer’s disease and related disorders.^11,12^

Available therapies against type-II diabetes include subcutaneous mealtime injections of pramlintide, an amylin analog, that reduces glucose concentrations by regulating the plasma glucagon level, slowing down gastric emptying, and improving satiety. ^13^ Unlike amylin itself, pramlintide resists fibril formation and inhibits or disaggregates amylin aggregates. However, because of the short half-life time of pramlintide (~ 50 minutes), the drug needs to be administered frequently, leading to undesired side-effects such as nausea and hypoglycemia.^14^ Hence, there is still a need for the development of new drugs with amylin-mimetic properties that reduce less the life quality of patients.

Short half-life time and the need for frequent administration of the drug are common complications in peptide-based therapies. One way of circumventing these problems is the use of D-retro Inverso (DRI) peptides. DRI peptides are made up of D-amino acids, stereo-chemically mirror-images of the L-amino acids, and have at the same time their sequence of amino acids reversed. As a consequence, DRI peptides resemble the parent peptide and have similar biological activity, but are resistant to proteolytic digestion and therefore will have longer half-life times.^15^ Several studies have reported therapeutic effectiveness of DRI peptides, for instance, for the treatment of tumors.^16^ Note, however, that the symmetry is not complete and that there are subtle structural variations between DRI-peptides and their L-parents. These differences can impart new chemical properties and might even enhance the potency of the peptides as drugs. For instance, Daggett and co-workers showed that peptides made of alternating L and D amino acids bind preferentially to toxic oligomers instead of monomeric and fibril forms of A*β*, inhibiting A*β* aggregation and reducing cytotoxicity in mouse and *C. elegans* models of Alzheimer’s disease.^17^

The above mentioned studies hint at the possibility that DRI-amylin may provide an alternative to pramlintide as a drug for targeting type-II diabetes. Hence, in this paper, using all-atom molecular dynamics simulations, we first study the ability of DRI amylin to form amyloids and then explore how the presence of DRI amylin alters the elongation and the stability of L-amylin fibrils. We find that DRI-amylin differs from L-amylin by the twist of *β*-sheets, but has similar or only slightly lower stability, resulting from a rearrangement of contacts and loss of hydrogen bonds. While DRI-peptides interfere with elongation, their main effect is lowering the stability of hybrid amylin fibrils by easing separation of protofibrils, suggesting a potential for use of DRI-amylin as inhibitor of amylin fibril-formation, Our investigations also indicate that insertion of DRI-proteins in L-assemblies can be used as a tool to probe the role of critical hydrogen bonds in supramolecular assemblies or aggregates. In the present study allowed us such insertion to identify the crucial role of Asn ladders and Ser-Ser bifurcated hydrogen bonds for stabilizing amylin fibrils.

## 2 Materials and methods

### 2.1 Model construction & simulation details

In order to connect our present study with earlier work investigating the stability of L-amylin aggregates,^12^ we have first constructed L-amylin decamers from U-shaped chains of a model provided by the Tycko’s lab.^18^ Each of the ten chains forms a *β*-strand-loop-*β*-strand motif, where the two *β*-strands are made up of residues 8–17 and 28–37, and the loop region is formed by residues 18–27. The decamer is composed of two pentamers (called by us “protofibrils”) with in each of the five “layers” two anti-parallel chains packed together at the C-terminal ends by polar interactions, hydrogen bonds involving Ser29 and Asn31; and hydrophobic interactions between Ala25–Asn35 and Leu27–Gly33. A representation of our model is shown in Figure 1.

**Figure 1:**
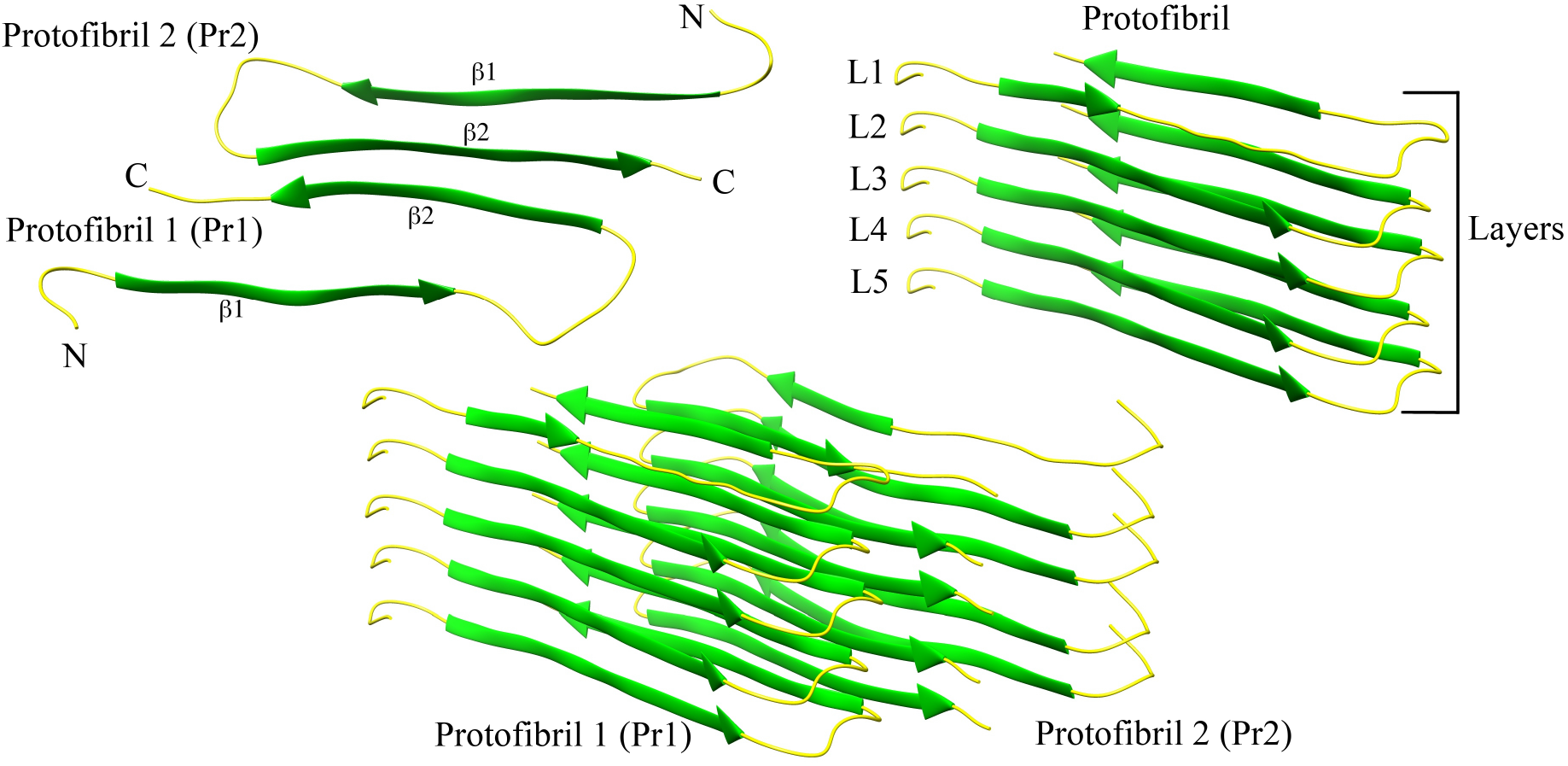
A two-fold human amylin fibril fragment. Chains in each layer interact by the anti-parallel C-terminal ends (a). Each protofibril is a stack of parallel chains (b). The arrangement of the two protofibrils is shown in (c)

Using the protocol described in Xi & Hansmann, ^19^ we have constructed a DRI version of this decamer by replacing the backbone atoms of amylin as such: nitrogen (N) to carbon (C), C to N, hydrogen (H) to oxygen (O), and O to HN (a hydrogen bound to a nitrogen). The resulting structure is refined by energy minimization and a subsequent short molecular dynamics simulations. The resulting model is shown in Suppl. Figure 1 and has by construction the same overall structure than the L-amylin decamer. However, each of the chains is now made of D-amino acids, with the sequence of residues inverted. Because of this inversion, and in order to compare more easily L and DRI amylin chains, we index the residues in DRI-amylin starting from the C-terminus instead (as in the case of L-amylin) in the usual order starting from the N-terminus.

Various hybrid models were generated by applying the above procedure only to a subset of pre-selected chains, leading to fibrils with a mixture of L and DRI peptides. These hybrid models are summarized in Figure 2 and probe possible scenarios by which DRI-amylin chains may interact with L-fibrils. For instance, the fibril model (4L-1D)*2 describes an elongation of a two-layer octamer ((4L)*2) by a DRI-peptide in each protofibril. Note that in the above notation, we abbreviate the ‘DRI’ by a ‘D’. In a similar way describes the model (2L-1D-2L)*2, a case where a DRI-peptide has been incorporated in each of the protofibrils. Cross fibrillization between L and DRI amylin is probed with assembly geometries (L-D-L-D-L)*2, abbreviated as (L/D)*2, and (L-D-L-D-L)*(D-L-D-L-D), abbreviated further as (L/D)*(D/L). Finally, the interaction between a DRI amylin pentamer with a L-amylin pentamer is probed by the fibril geometry 5D*5L.

**Figure 2:**
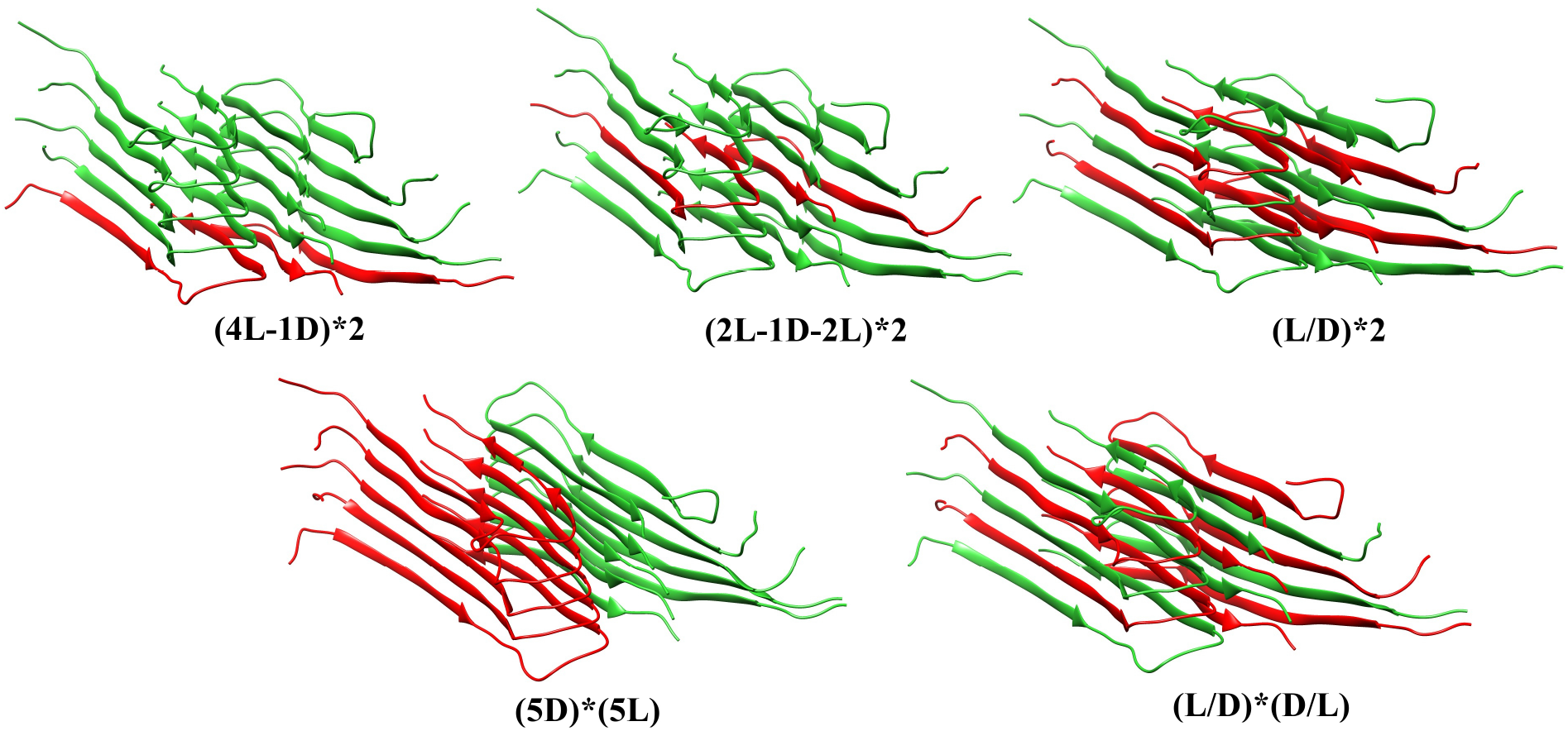
Various models of hybrid fibrils builds from mixtures of L and DRI amylin chains

We use for our molecular dynamics simulations the software package GROMACS 2018.1,^20,21^with the protein interactions approximated by the CHARMM36 force field.^22^However, in order to compare our results with our earlier work and to gauge any force field dependence of our results, we have also simulated the L-amylin fibril fragment using the AMBER (ff99SB) force field.^23^For each run, we place the respective decamer in the center of a cubic box with 12 Å distance to the edges and fill the box with TIP3P water.^24^ Because of periodic boundary conditions, we evaluate electrostatic interactions by particle-Ewald summation ^25,26^and a cut-off of 1.2 *nm* is used for calculation vdW-interactions.The resulting systems are energy-minimized by steepest descent, followed by short (500 *ps*) molecular dynamics in an NVT ensemble, and subsequent 500 *ps* in an NPT ensemble. Temperature and pressure are controlled by a Parrinello-Danadio-Bussi thermostat^27^and Parrinello-Rahman barostat,^28^ and are set to T=310 *K* and 1 *bar*. The integration step is 2 *fs*, For each system, we follow three independent trajectories (starting from different velocity distributions) over 200 *ns*.

We analyze the resulting trajectories using the tools provided by the GROMACS package. One important quantity that we measure is the root-mean-square deviation of C*α*-atoms (RMSD) with respect to the corresponding start configurations, excluding the flexible first seven residues. Monitoring the time evolution of the RMSD for the various models, we find that our simulations converge after 120 *ns* (Suppl. Figure 2). Therefore, we consider only the last 80 *ns* of the 200 *ns* long trajectories for further analysis (*i.e.*, for calculation of average properties). Other quantities which we measure and analyze include the radius of gyration (Rg), solvent-accessible-surface-area (SASA), root-mean-square-fluctuation (RMSF), hydrogen bonds, dihedral angles, twist angle, hydrophobic contacts and CC–interface contact distance. Hydrogen bonds are defined by a distance cut-off of 3.5 Å between the donor & acceptor atom and an angle cut-off of 30°. Similarly, a hydrophobic contact is defined by the condition that the distance between two residues (*i* and *j*, with |*i* − *j*| > 3) is less than 4.5 Å. We define the CC interface contact distance as the distance between the C*α* atoms of the residue pairs Leu27–Gly33, Ser29–Asn31, Asn31–Ser29 and Gly33–Leu27. Facing each other, these residues contribute towards fibril packing by forming hydrogen bonds and hydrophobic or polar contacts. We further calculate for the penultimate chains of a protofibril the vector between the C*α*-atom of Gln10 and the C*α*-atom of Leu16. We define the twist angle for the *β*1-strand in a protofibril by the angle between these two vectors. In a similar way, we define the twist angle for the *β*2-strand by the corresponding vectors that point from the C*α*-atom of residue Leu27 to the C*α*-atom of Gly33.

## 3 Results and discussion

### 3.1 Force field dependence on amylin fibril simulations

A few years ago, we have studied in our lab the stability of wild type and various mutations of amylin fibril fragments.^12^ The simulation of these decamers (built from L-amino acids) relied on the AMBER (ff99SB) force field. Hence, in order to connect with our earlier work, we have started our investigation by first simulating our L-amylin fibril model (also a decamer) using the AMBER ff99SB force field. As our new results agree within the error bars with previous work,^12^see Suppl. Figure 3, we conclude that our present simulation set-up is correct. However, we prefer to replace in the present work the previously used AMBER ff99SB by the newer CHARMM36 force field,^22^which in our experience is more suitable for simulations of intrinsically disordered and aggregated proteins. Contrasting the ff99SB simulations of the L-amylin fibril with corresponding simulations relying on CHARMM36 allows us also to estimate the force field dependence of our results; and important factor to consider when comparing the stability of L-amylin and DRI-amylin aggregates. For this purpose, we have calculated the distribution of various quantities measured in the two sets of simulations. These distributions are shown in Figure 3. While the distribution of the radius of gyration (Rg), a measure for the compactness of configurations, is similar for the two sets, the situation is different for root-mean-square deviations (RMSD) to the respective start configurations. For this quantity, one derives a much broader distribution from the CHARMM trajectories than from the AMBER trajectories. This broadening goes together with a shift of the solvent-accessible-surface-area (SASA) values toward more solvated configurations in the CHARMM36 simulations. Together with the raised root-mean-square-fluctuations (RMSF) for each residue, suggesting a more diverse ensemble of configurations in the CHARMM36 simulations, these changes illustrate the improvements in force field development of the last years: modern force fields are less focused on the folded states as they aim to describe also correctly the energetics of unfolded configurations.

**Figure 3:**
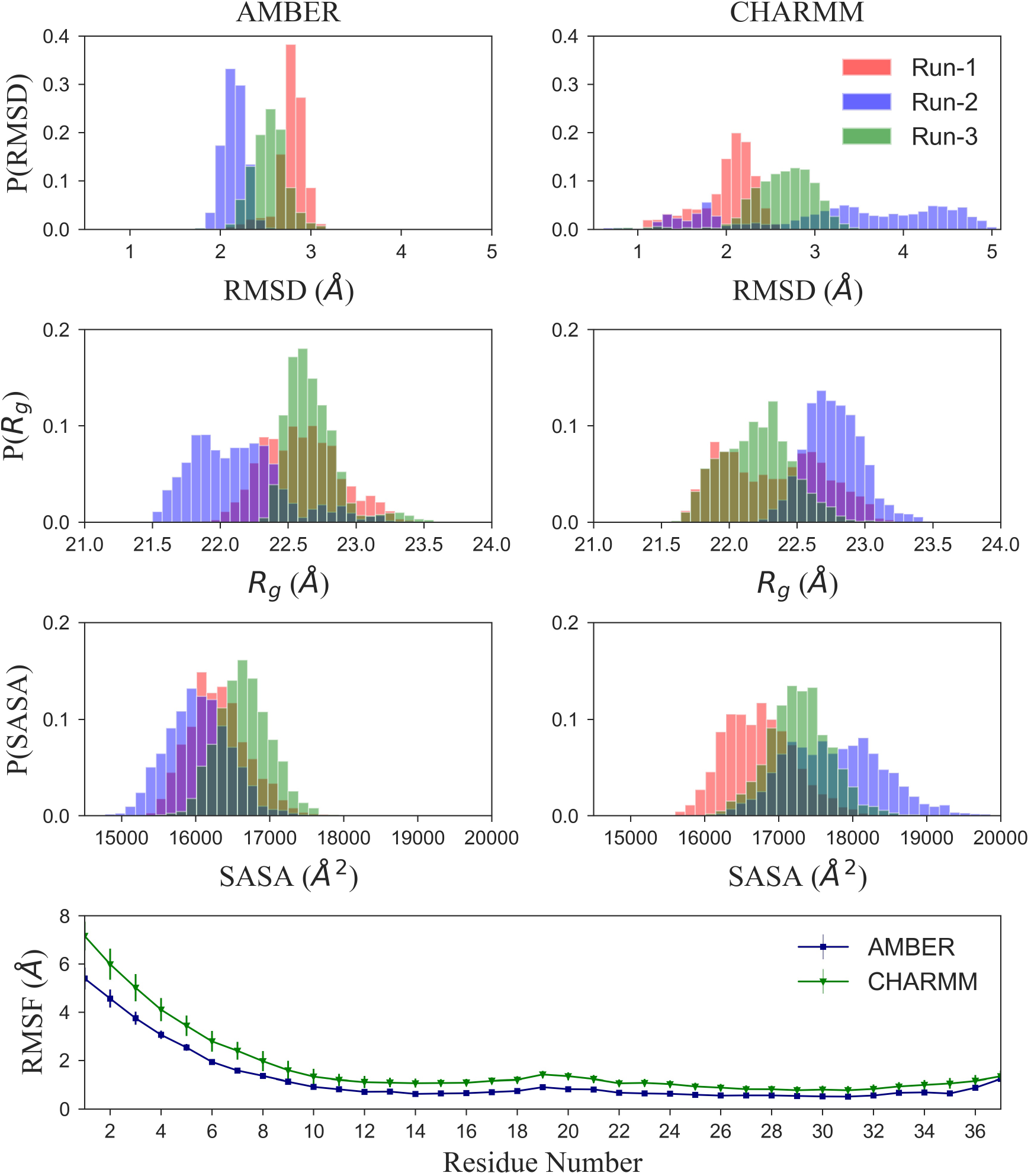
Comparison of molecular dynamics simulations of L-amylin fibril fragment using either AMBER ff99SB or CHARMM36. For this purpose, we show in (a) the normalized distribution of the root-mean-square deviation (RMSD), (b) the radius of Gyration (Rg), and in (c) of the solvent-accessible-surface-area (SASA). In (d), we show the root-mean-square-fluctuation (RMSF) of the C*α* atoms. Values are averaged over all chains in the fibril fragment and all three trajectories.

### 3.2 Comparison of L-amylin and DRI-amylin fibrils

While the structural similarity between DRI-peptides and their L-parents often results in similar biological properties and activity, the existing subtle differences may alter amyloid formation and stability. For instance, in an earlier study, we found noticeable deviations in the stability of L and DRI A*β*40 and A*β*42 peptide fibrils, suggesting that DRI A*β* peptides enhance fibril formation in hybrid systems. However, it is not clear whether these earlier results can be generalized to amylin; and for this reason, we compare here first the stability of L-amylin and DRI-amylin fibrils.

In our earlier work, we could connect some of the stability disparity between L and DRI A*β*40-fibrils to the dissimilar twist angles seen in the arrangements of the U-shaped chains. For this reason, we have measured here in both protofibrils the twist angles for the *β*-strands *β*1 and *β*2. While the Ramachandran plots in Suppl. Figure 4 indicate that the sampling of the backbone *ϕ*/*ψ* angles of DRI-amylin mirrors those sampled by L-amylin, and the magnitude of the twist angle varies little between DRI-amylin and L-amylin for the N-terminal *β*1-strands (L-amylin: −15.1° ± 6.6°, DRI-amylin: 12.8° ± 3.6°), the measured twist angles differ not only by their sign but also in magnitude for the C-terminal *β*2-strands. Here, the DRI twist angle is more than twice as large than the L-value (L-amylin: −3.3° ± 2.5°, DRI-amylin: 7.6° ± 3.8°) (Suppl. Table 1). Because of these dissimilar twist angles, the residue-residue distances at the CC-interface (dominated by polar residues) between the two protofibrils differ slightly (see Supple. table 2), resulting in a tighter packing at the CC interface in L-amylin fibril fragments than found in the DRI-amylin fibril model. However, the resulting loss in stability is small: the average root-mean-square deviation (RMSD), radius of gyration (Rg), solvent-accessible-surface-area (SASA) and RMSF as listed in Table 1 and Suppl. Figure 5 are similar or only slightly higher for DRI-amylin than for L-amylin fibrils; and no obvious differences are seen, when comparing individual trajectories (Suppl. Table 3).

**Table 1:**
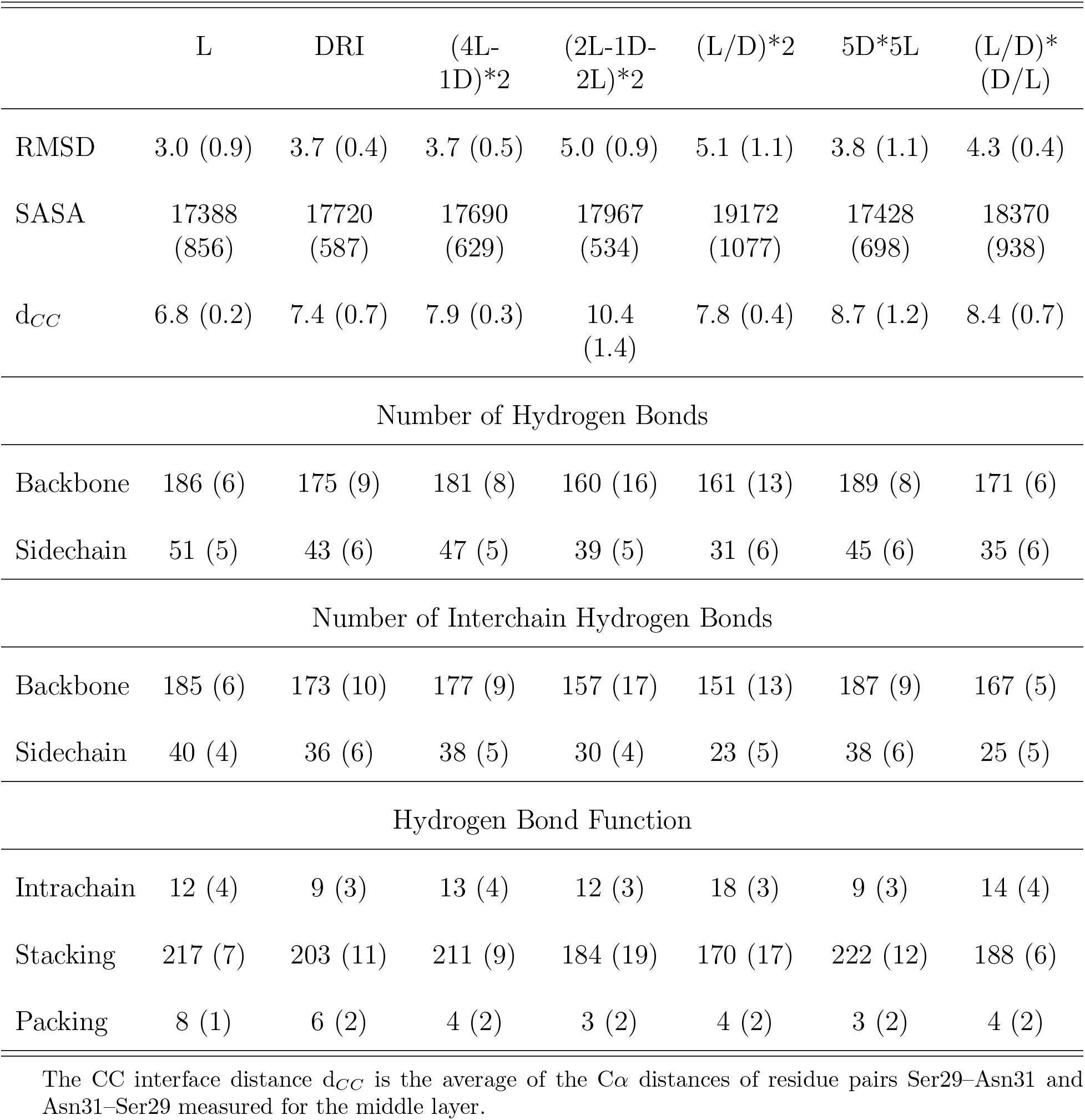
Structural changes in L-amylin (L), DRI-amylin (DRI) and Hybrid Amylin models in terms of average root-mean-square deviation (RMSD, in Å), average solvent-accessible-surface-area (SASA, in Å^2^), number of mainchain hydrogen bonds, sidechain hydrogen bonds, number of hydrogen bonds involved in stacking and packing, and CC interface distance d_*CC*_ (Å).

As the difference in the magnitude of twist angles is seen solely for the C-terminal *β*2-strands, the difference likely results from the interaction between chains on opposite protofibrils. The end of the N-terminal *β*-strand and the loop region connecting the two *β*-strands in each chain forms a hydrophobic core that contributes to the stability of the *β*-hairpin (*β*-strand-loop-*β*-stand motif) and to the stacking of layers. While the total number of these hydrophobic contacts differs little between L-amylin (193 ± 1) and DRI-amylin fibrils (192 ± 7), rearrangements are observed in the inter-strand hydrophobic contacts for the residues Leu12, Ala13, Phe15, Val17, Ala25, Ile26, Leu27 and Val32, with 67 contacts seen only for L-amylin, and 68 only for DRI amylin. For instance, in L-amylin, residue Phe15 on strand *i* makes a hydrophobic contact with Ala13 on strand *i* + 1, and Val17 on strand *i* makes another hydrophobic contact with Phe15 on strand *i* + 1. In DRI-amylin, the residues switch places, *i.e.*, it is now the Ala13 on strand *i* that forms a contact with the Phe15 on strand *i* + 1, while the Phe15 of strand *i* is now paired with the Val17 of strand *i* + 1. Similar re-arrangements of hydrophobic contacts are observed for residues Leu12, Ala25, Ile26, Leu27 and Val32. Only present in L-amylin is a hydrophobic contact between residue Leu12 on strand *i* and Ala13 on the neighboring strand *i* + 1, while DRI-amylin has a contact between Ala8–Ala8 on neighboring strands that is not found in L-amylin. As more than 60 % of the hydrophobic contacts are preserved in both forms, and lost contacts are replaced by newly formed, it appears that hydrophobic contacts add in similar way to the stability of DRI-fibrils and L-fibrils.

With the exception of an intrachain sidechain hydrogen bond connecting residues Asn35 with Tyr37, and a transiently formed mainchain hydrogen bond connecting N and C-termini, are all hydrogen bonds interchain bonds, and therefore essential for fibril stability. Mainchain hydrogen bonds link *β*-strands of stacked chains in the same protofibril, while sidechain hydrogen bonds also contribute towards the packing of the two protofibrils. DRI-amylin has 173 ± 10 backbone interchain hydrogen bonds and 36 ± 6 sidechain interchain hydrogen bonds, approximately twelve mainchain and four sidechain hydrogen less than L-amylin (185 ± 6 mainchain hydrogen bonds and 40 ± 4 sidechain hydrogen bonds), see Table 1. While donor and acceptor atoms are switched, DRI-amylin preserves most mainchain hydrogen bonds found in L-amylin fibrils. The exception, leading to the unequal number of mainchain hydrogen bonds, are two recurring inter-layer hydrogen bonds which in L-amylin are formed between residues bordering the *β*-turn region but are in DRI-amylin fibrils only seen with a much lower frequency. The first of these hydrogen bonds connects residue Gly24 of one layer with Ala25 of the layer above and appears in L-amylin with a frequency of (90 ± 10) %, but in DRI-amylin only with a frequency of (13 ± 4) %. Gly24 and Ala25 are part of the large hydrophobic core region of amylin that stabilizes the fibril by hiding the core residues from water. The second L-amylin-specific hydrogen bond connects in a similar fashion residue on neighboring layers, however, the two residues, His18 and Ser19, are on the opposite side of the *β*-turn region. This hydrogen bond is found in L-amylin in (74 ± 28) % of configurations, but only in (20 ± 2 %) of DRI-amylin configurations. The difference in frequency with which these in total 16 inter-layer hydrogen bonds are observed leads to an effective loss of eleven *β*-turn stabilizing hydrogen bonds in DRI-amylin fibrils, contributing to the slightly lower structural stability of DRI-amylin.

The difference in the number of sidechain hydrogen bonds measured in L and DRI fibrils is surprising as DRI peptides preserve the sidechain geometry of their parent L-peptide. Both fibrils have Asn14–Asn14 hydrogen bonds, and in most cases Asn22–Asn22 hydrogen bonds (which otherwise are replaced in DRI-amylin fibrils by His18–Asn22 hydrogen bonds), that both connect chains on subsequent layers of the same protofibril. However, visual inspection shows that the orientations of the sidechains of Ser29 and Asn31 differ in the two fibrils (Figure 4(d)). In L-amylin, the sidechains of Asn31 residues are aligned on top of each other (homo-stacking), forming Asn31–Asn31 sidechain hydrogen bonds (Figure 4(b)). On the other hand, the Ser29 residues from chains in opposite protofibrils face each other, allowing the formation of Ser29–Ser29 sidechain hydrogen bonds (Figure 4(a)). This is different in DRI-amylin where Ser29 can now only form hydrogen bonds with Asn31 of the opposite protofibril (either of the same or a neighboring layer) (Figure 4(c)). It is this re-arrangement of hydrogen bonds between the sidechains of residues Ser29 and Asn31 in neighboring chains that leads to the lower number of sidechain hydrogen bonds in the DRI-fibril.

**Figure 4:**
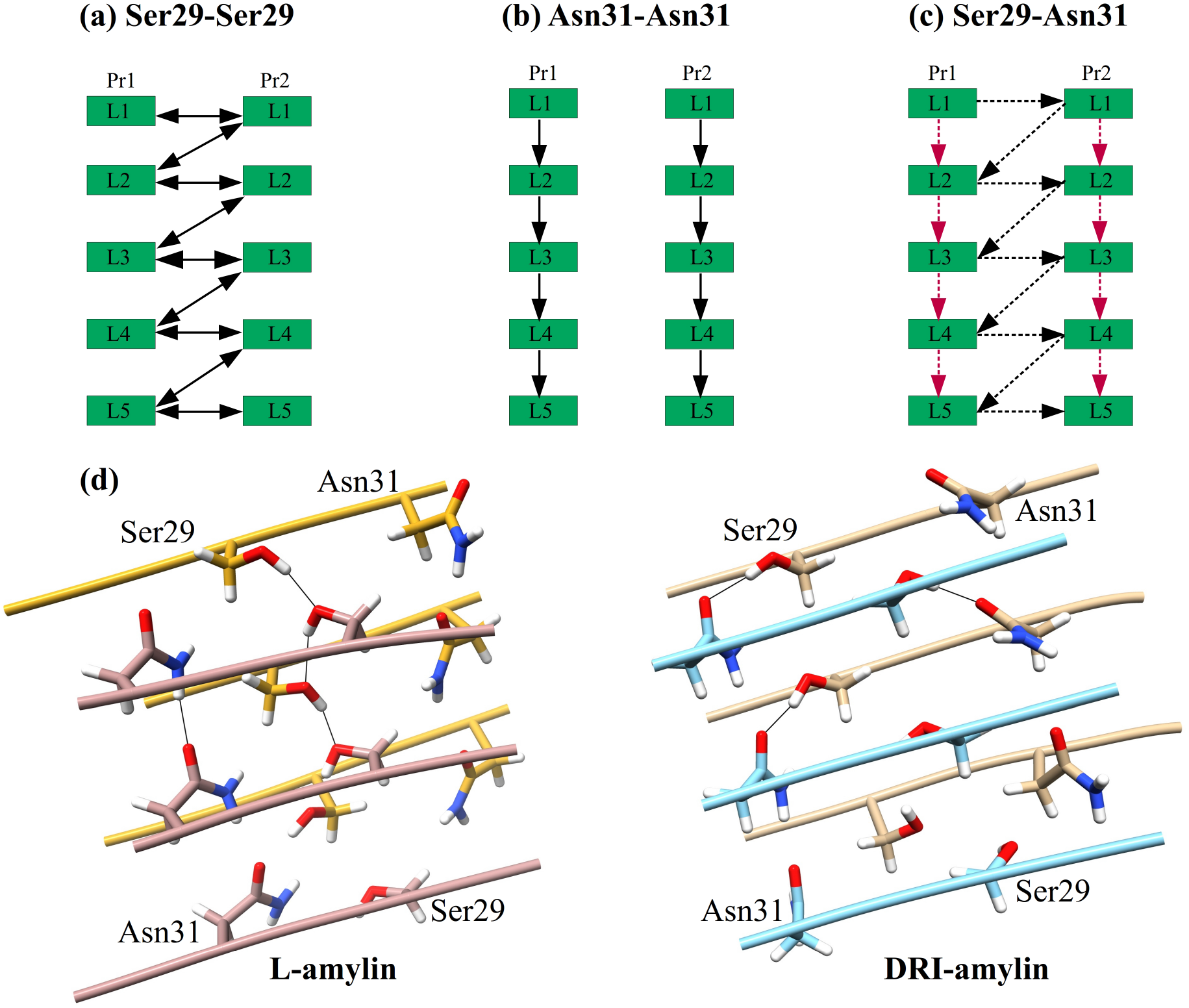
Schematic of the interchain sidechain hydrogen bond observed in L-amylin and DRI-amylin. In L-amylin, Ser29 forms a hydrogen bond with neighboring Ser29 (Ser29@OG1–Ser29@HG1) of the opposite protofibril (a), and Asn31 forms a hydrogen bond with Asn31 (Asn31@OD1–Asn31@2HD2) on subsequent layer of the same protofibril (b) However, in DRI-amylin, Ser29-Ser29 hydrogen bonds do not exist and on average half of the Asn31-Asn31 hydrogen bonds (drawn in red) are replaced by hydrogen bonds between Asn31 and Ser29 (drawn in black) (c). This is because the sidechain of Ser29 and Asn31 are flipped as can be seen in (d). Arrows point from donor to acceptor, and competing hydrogen bonds are drawn with dotted lines.

Since in L-amylin the interface between the two protofibrils is formed by the (anti-parallel) C-terminal strands of each chain, the sidechain of residue Ser29 on a given chain can form two hydrogen bonds with the sidechains of Ser29 residues located on chains in the other protofibrils. One hydrogen bond connects Ser29 residues on the same layer and is seen in about (78 ± 23 %) of all configurations. The other hydrogen bond connects Ser29 residues on neighboring layers *i* and *i* − 1 (but different protofibrils) and is seen in (88 ± 19) % of configurations. The average distance between the donor-acceptor pair is in these two hydrogen bonds comparable: (2.80 ± 0.04) Å (when connecting residues located on the same layer) and (2.79 ± 0.03) Å (when connecting residues on staggered layers). Note that these two hydrogen bonds are bifurcating hydrogen bonds, with the donor and acceptor atoms (OG1 and HG1) switching with a period of (2.2–2.3) *ps* between the two hydrogen bond forming residues (Data not shown). As a consequence, the Ser29–Ser29 hydrogen bonds support in L-amylin fibrils both the stacking of chains in each protofibril, and the packing of the protofibrils. This is not the case in DRI-amylin fibrils where these bifurcating hydrogen bonds are not observed (the corresponding frequencies are (5 ± 11) % and (5 ± 6) %) for both types.

This effect is partially compensated by alternative hydrogen bonds that Asn31 can form in DRI-fibrils but not in L-fibrils, where Asn31 can only form a single hydrogen bond with an Asn31 on a chain in the above layers of the same protofibril. These hydrogen bonds appear with a frequency of (87 ± 16) % of configurations and stabilize the stacking of chains in each protofibril. The same kind of hydrogen bond is also seen in DRI-amylin fibril but only with a frequency of (42 ± 27) %, *i.e.*, with half the probability found in L-amylin, and therefore reducing further the cohesion between layers in the protofibrils. However, when in in DRI-amylin fibrils not forming an Asn31–Asn31 hydrogen bond, the Asn31 residue on a given chain can form instead a hydrogen bond with a Ser29 on a chain located either on the same layer or on a neighboring layer of the opposite protofibril. These nine possible Asn31–Ser29 hydrogen bonds are observed in (51 ± 35) % of all DRI-fibril conformations, but only in less than 2 % of L-amylin fibrils. While the donor-acceptor distance in these particular hydrogen bonds is with (2.77 ± 0.02) Å comparable to the of Ser29–Ser29 hydrogen bond found in L-amylin, their lifetime is with ~1.3 *ps* (Data not shown) very short, and they are not bifurcating hydrogen bonds as the donor is always Ser29 and acceptor is always Asn31. Nevertheless, replacing in DRI-fibrils the Ser29–Ser29 hydrogen bonds seen in L-amylin, the Asn31–Ser29 bonds stabilize in a similar way the packing of the protofibrils.

However, the loss of all of the nine Ser29-Ser29 hydrogen bonds and on average half of the eight Asn31–Asn31 hydrogen bonds is only partially compensated by the formation of on average half of the nine Asn31–Ser29 hydrogen bonds possible in the DRI-amylin fibril fragment. While on average in L-amylin one Asn31–Asn31 sidechain hydrogen bond connects two chains located on layers of the same protofibril, the number is reduced by half in DRI-amylin. Similarly, while in L-amylin on average one Ser29-Ser29 hydrogen bond per chains connects the two protofibrils, only half of these are replaced in DRI-amylin by Asn31–Ser29 hydrogen bonds. Hence, the net-effect of the diverse hydrogen bond pattern is a reduction of stability in DRI amylin fibrils by four sidechain hydrogen bonds.

Our discussion of the differences in hydrogen bonding also explains the rather small differences in stability between L-amylin fibrils and DRI-fibrils. In L-amylin fibrils are neighbouring layers hold together by on average 23 mainchain hydrogen bonds and four sidechain hydrogen bonds per pair of chains, leading to a total of 217 (7) hydrogen bonds that stabilize stacking of chains. In DRI-amylin are the corresponding numbers 21 mainchain hydrogen bonds and three sidechain hydrogen bond per pair of chains, leading to a total of 203 (11) hydrogen bonds that stabilize stacking of chains. Hence, stacking of chains is less stable in DRI amylin fibrils by about two hydrogen bonds per pair of chain, a total reduction in stacking of about 7 %. On the other hand, the two protofibrils are in the L-amylin fibril hold together by on average eight sidechain hydrogen bonds, but only by six bonds in the DRI-amylin fibril, reducing the packing stability by about 25 %.

While the loss of hydrogen bonds in DRI-amylin fibrils reduces only slightly the overall stability, and individual *β*-strands keep their structure, it still leads to notable structural differences. The looser packing of the individual protofibrils, resulting from the smaller number of sidechain hydrogen bonds, leads to a more twisted conformation, with the DRI twist angle for the C-terminal *β*2-strands twice as large as the corresponding L-value. In Lamylin is the twisting of the C-terminal *β*2-strands restrained by the hydrogen bonds formed between the side chains of the Ser29 residues on the two protofibrils (Ser29-Ser29), leading to twist angles of −2.9° ± 0.5° and −3.6° ± 0.4° (see Suppl. Table 1). The effective loss of these hydrogen bonds in DRI-amylin is only partially compensated by the newly formed sidechain hydrogen bonds between Ser29 and Asn31, but not sufficiently to counterbalance the intrinsic propensity of the protofibrils to twist. As a consequence, we observe a higher twist angle in the *β*2-strands of the protofibrils of DRI-amylin (7.2°± 2.9° and 8.0°± 4.5°), see Suppl. Table 1. The consequence of this larger twist are the two protofibrils less close than in L-amylin. For instance, we measure as CC-interface distance a value of (7.4 ± 0.7) Å, compared to (6.8 ± 0.2) Å for L-amylin fibrils.

### 3.3 Stability of Hybrid Assemblies

Despite the only marginal structural differences, there are distinct differences in the hydrogen bond pattern between L-amylin fibrils and such made of DRI-amylin, with the effect most pronounced for sidechain hydrogen bonds that can alternate (or oscillate) between two residue pairs sharing one residue as either donor or acceptor (in our case Ser29). This suggests that insertion of DRI-proteins in L-assemblies may be an alternative to mutations for probing the role of such hydrogen bonds in supra-molecular assemblies or aggregates. Another possible application would be to use these differences to modulate formation and growth of fibrils.

Fibril formation is a nucleation process where rapid fibril growth follows the formation of a critical nucleus. The long lag-phase required for nucleation can be shortened by orders of magnitude through seeding, *i.e.*, the addition of preformed fibrils as seeds. These seeds do not have to be of the same protein, as amyloids can be also formed by cross-seeding with other proteins.^12^The latter process is not fully understood, and is especially interesting for mixtures of L and DRI peptides where the differences in chirality impart subtle differences in the hydrogen bond and hydrophobic interactions (as seen in the previous section) that can either promote or inhibit seeding. Hence, in order to understand how the presence of DRI-amylin affects growth and stability of L-amylin fibrils we have studied the stability of various hybrid constructs made up of L and DRI amylin.

In the first set of simulations, we have looked into the effect of DRI-peptides on the elongation of amylin fibrils. For this purpose, we considered the case of DRI-peptides attached at the end of both layers of a double-layer L-amylin fibril, *i.e.*, the fibril construct (4L-1D)*2. This decamer models the elongation of a double-layer L-amylin octamer ((4L)*2) by a DRI-peptide at each layer. As can be seen from Table 1, the (4L-1D)*2 fibril has 0.7 Å higher average RMSD (3.7 ± 0.5) Å, 0.3 Å higher Rg (22.7 ± 0.4) Å and 302 Å^2^ more SASA (17690 ± 629) Å^2^ values than the corresponding L-amylin decamer, and therefore is only slightly less stable. Since hydrogen bonds across the chains are the major player in the elongation process of the fibril assembly, we have calculated the number of mainchain and sidechain hydrogen bonds, and observe that most of the mainchain (181 ± 8) and sidechain hydrogen bonds (47 ± 5) are conserved in (4L-1D)*2 hybrid fibril, with discrepancies only at the L and DRI interface. In a pure L-fibril, or in a pure DRI-fibril, each *i* residue in one chain forms two mainchain hydrogen bonds with the residues in the subsequent layer: one with *i* − 1 residue with the backbone amide hydrogen and the other with *i*+1 residue with the backbone carbonyl oxygen (Figure 5). However, at the L–DRI interface, the *i* residue from L-chain forms both mainchain hydrogen bonds with the *i* − 1 residue from DRI-chain, *i.e.*, the DRI strand is shifted in relation to the rest of the fibril by one residue towards the terminus. As a consequence, the loop region and C-terminus end of the DRI-chain are no longer in close proximity to the loop region and the C-terminus end of the underlying L-chain, hindering the formation of mainchain hydrogen bonds in this region, specifically between the residue pairs Asn22–Phe23 (Pr2L4–Pr2L5), Ala25–Ile26 (Pr1L4–Pr1L5) & Gly24–Ala25 (Pr1L4–Pr1L5) (beta turn region) and Ser34–Asn35 (Pr2L4–Pr2L5) & Thr36–Tyr37 (Pr2L4–Pr2L5) (terminal regions). The hydrogen bonds between the residue pairs Asn22–Phe23, Gly24–Ala25 & Ala25–Ile26 occur with a frequency of (91 ± 19) %, (95 ± 5) % and (98 ± 1) % at the interacting L–L interface, while the corresponding values for the L–DRI interface are within the error bars consistent with zero. The loss of four backbone hydrogen bonds leads to a weakening of the inter-layer stacking at the interface between L and DRI chains, causing in turn re-arrangement of hydrophobic contacts at this interface residues Leu12, Ala25, Ile26, Leu27 and Val32.

**Figure 5:**
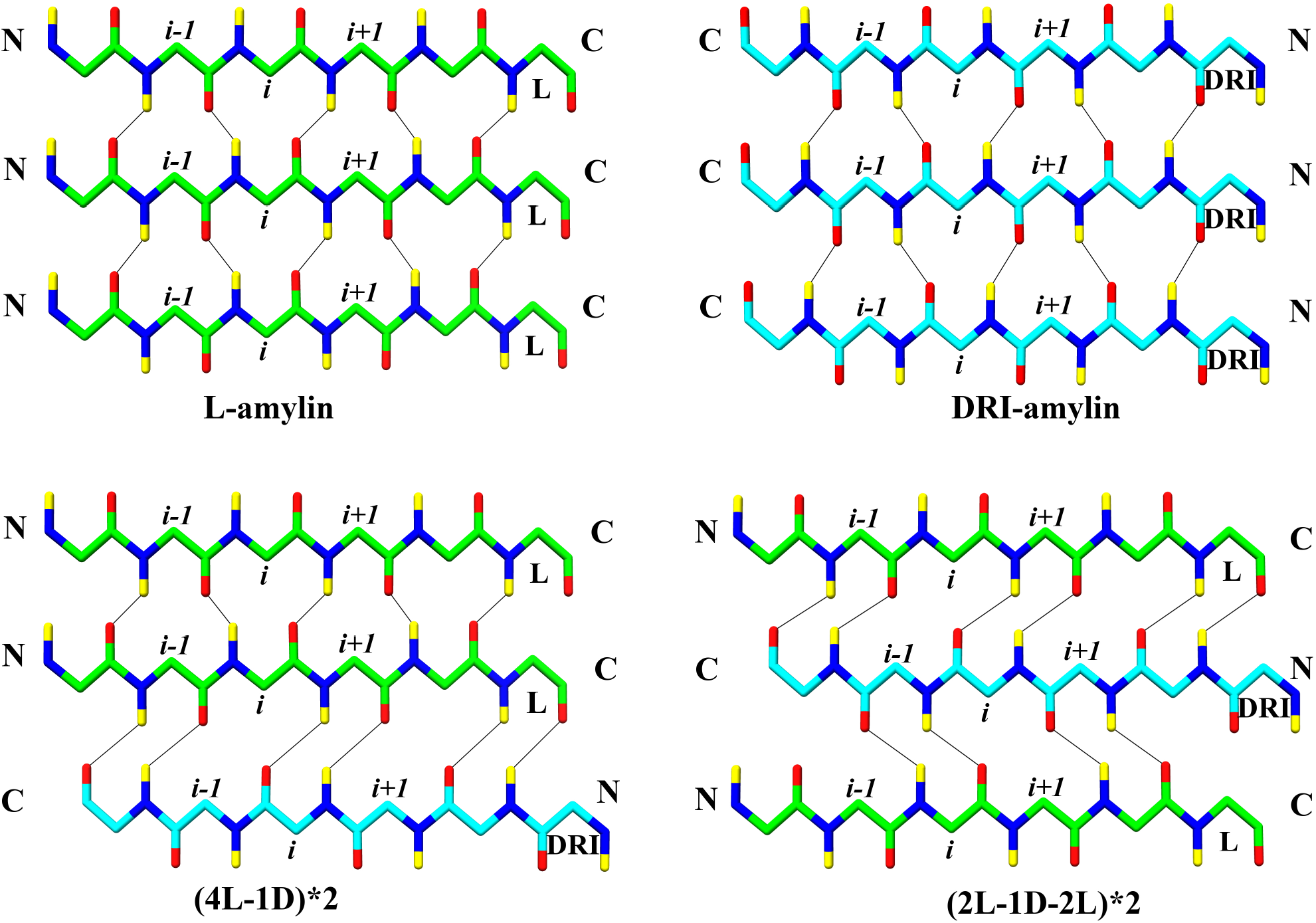
Interchain hydrogen bonds formed in L-amylin, DRI-amylin and hybrid fibrils. Between L–L and DRI–DRI strands, interchain mainchain hydrogen bonds are formed connecting the *i* residue from one strand to the *i* − 1 and *i* + 1 residue from another strand. In case of a L–DRI interface, the *i* residue from one strand forms interchain mainchain hydrogen bonds with the *i* − 1 residue or with the *i* + 1 residue from another strand. Only backbone atoms are shown for clarity. Oxygen atoms are coloured in red, nitrogen in blue, hydrogens in yellow and C*α* and C atom of L-chain is coloured in light green, while that of DRI-chain is coloured in cyan. Note that residues are counted from the C-terminal in DRI chains.

Since the DRI-amylin chain is shifted by one residue in relation to the L-amylin chains, we expect to see also differences in the sidechain hydrogen bond pattern at the L–DRI interface. For instance, the Asn31–Asn31 hydrogen bond is lost at the interface between L and DRI chains (Pr1L4–Pr1L5 and Pr2L4–Pr2L5). However, the attachment of DRI at the end of the L-amylin fibril affects also the network of Ser29–Ser29 hydrogen bonds beyond the L and DRI interface where these hydrogen bonds now appear only with a frequency of (20 ± 20) %. In pure DRI fibrils, the loss of Ser29–Ser29 hydrogen bonds is partially compensated by the formation of hydrogen bonds between Ser29 and Asn31, but this is not the case in the (4L-1D)*2 model. Since the Asn31–Asn31 side chain hydrogen bonds are still retained in the L–L layers, though appearing with a lower frequency of (45 ± 36) %, the hydrogen bond forming atoms of Asn31 are not available to form hydrogen bonds with Ser29 to make up for the loss of Ser29–Ser29 hydrogen bonds. As a consequence, there are at the L-DRI interfaces between layers 4 and 5 in the two protofibrils three sidechain hydrogen bonds less involved in stacking, approximately one hydrogen bond at each L-DRI interface. On the other hand, the two L-DRI intetfaces between chains located on opposite protofibrils, lead to the loss of four sidechain hydrogen bonds for packing, two for each L-DRI interface. The loss of packing-supporting hydrogen bonds is because the larger twist of the C-terminal *β*2-strand in the DRI-amylin chain affects also the corresponding strands of the L-amylin chains in the other layers of the fibril which have about three times larger twist angles than seen in pure L-amylin fibrils. Hence, the addition of the DRI-fibril leads to the loss of the sidechain hydrogen bonds that stabilize the packing of the two protofibrils: only four hydrogen bonds stabilize the packing, while in L-amylin fibrils the packing is stabilized by eight hydrogen bonds, and by six hydrogen bonds in DRI-amylin fibrils. Hence, elongation of a L-fibril by DRI peptide is energetically less favorable than elongation by an L peptide, since the packing of layers is reduced at the interface by four mainchain hydrogen bonds, and the packing between protofibrils by four hydrogen bonds, reducing the interaction between the protofibrils even beyond the interfacing layer (see Table 1). As a consequence is the CC interface distance with 7.9 Å substantially larger than in either L– or DRI amylin fibrils.

While the simulations of the fibril construct (4L-1D)*2 allows us to probe the elongation of existing L-fibrils by DRI-amylin, and the nucleation of L-amylin fibrils by DRI-amylin seeds, they describe settings where there is only one interface between L and DRI peptide. Hence, these simulations cannot tell us about the stability of hybrid fibrils where DRI peptides are incorporated within L-layers, *i.e.*, whether DRI-amylin as a fibril breaker or enhances the stability of amylin fibrils. The simplest system to study this question is an arrangement (2L-1D-2L)*2, where in each protofibril the central chain is a DRI-amylin, surrounded by two L-amylin chains on each side. We find that this hybrid construct is even less stable than the (4L-1D)*2 construct, with a higher RMSD (5.0 ± 0.6 Å) and more solvated structures seen at the end of the trajectories (see Table 1). As a consequence, this model has 199 (17) hydrogen bonds, about 30–40 hydrogen bonds less than the L-amylin model (or even the (4L-1D)*2 model), and about 20 less than DRI amylin fibril (see Table 1). All of the lost hydrogen bonds are interchain. The differences in hydrogen bonding and hydrophobic contacts are again concentrated at the interfaces between L and DRI peptides. Since in (2L-1D-2L)*2, the DRI-chain interfaces with two L-amylin chains, the *i* residue from the DRI-chain forms both backbone hydrogen bonds either with the *i* + 1 residue of the L-chain in the next layer, or with the *i*+1 residue of the L-chain on the preceding layer (see Figure 5). However, at both interfaces are hydrogen bonds lost in the beta turn region (Asn22–Phe23 (47 ± 46 %), Gly24-Ala25 (8 ± 23 %) & Ala25-Ile26 (36 ± 44 %)) and terminal region (Ser34-Asn35 (0 %) & Thr36-Tyr37 (0 %)). In total, there are 157 (17) interchain backbone hydrogen bonds stabilizing the stacking of chains, 28 less than in L-amylin. Of these are 14 lost at the four L–DRI interfaces, *i.e.*, around four backbone hydrogen bonds per L–DRI interface, the same value as seen in the (4L-1D)*2 model. Note that while also 14 hydrogen bonds are lost at the four L–L interfaces, this loss results from increased fluctuation in the loop region and C-terminus due to inclusion of the DRI-chain, not from a relative shift of chains as in the case of L–DRI interfaces.

Similar to the (4L-1D)*2 case are in the (2L-1D-2L)*2 model the sidechain hydrogen bonds lost between Ser29–Ser29 and Asn31–Asn31 (14 ± 17) %, and only 3 (2) sidechain hydrogen bonds (less than one par pair of chains) contribute to the packing of the two protofibrils. However, in addition are also sidechain hydrogen bonds between Asn22–Asn22 (29 ± 35) % not compensated by formation of His18–Asn22 (3 ± 8) % sidechain hydrogen bonds. As a consequence, the stability of stacking in these fibrils is reduced by additional five sidechain hydrogen bonds when compared to pure L fibrils. Four of thee hydrogen bonds are lost at the four L–DRI interlayer interfaces, while three are lost at the two interlayer L-DRI interfaces in the (4L-1D)*2 model. Contributing to this loss of stacking stability is the lack of 28 hydrophobic contacts (when compared to L-amylin) involving residues Ala25, Ile26, Leu27 and Val32. Especially, the hydrophobic contacts between Leu27 & Val32 were lost across all the layers, while the contacts between residues Ala25 & Ile26 and Ile26 & Leu27 were lost only at the interfaces between L and DRI strands. The net effect of this loss of hydrophobic contacts and hydrogen bonds is a much larger reduction of stability than observed for pure DRI-amylin fibrils or (4L-1D)*2 construct, as seen, for instance, by the time evolution of RMSD (Suppl. Figure 2) or the much larger distance between the two protofibrils at the CC-interface (10.4 ± 1.4) Å, more than 4 Å larger than seen in the L-amylin fibril.

The fibril fragment (2L-1D-2L)*2 serves as a model for a defect in the fibril resulting from incorporating a DRI-chain into a L-fibril, a situation that may occur if the concentration of DRI-chains is small compared to that of L-chains. Once the concentration of DRI-chains is sufficiently large, alternating assemblies of L and DRI chains could appear instead if they were energetically favorable. We have considered three such assemblies. The first one is the model (L/D)*2, where in a given layer the chains are either both L or both DRI peptides. In the second model, 5D*5L, one protofibril is made of L-amylin chains and the other of DRI amylin. In the third model, (L/D)*(D/L), differ the chains in the two protofibrils, but also change in each layer. In neither case, do we see a stabilizing effect caused by the alternating L and DRI chains. In the (L/D)*2 model, designed by us for probing the effect of alternating L-and DRI amylin on the stacking of chains, we find that in comparison to the L-amylin fibril the average RMSD increases by two Å (5.1 ± 1.1 Å) and the average solvent accessible surface area (SASA) by 1784 Å^2^. Both the number of interchain backbone (151 ± 13) and sidechain hydrogen bonds (23 ± 5) is lower than in all cases, and only partially compensated by an increase in intrachain hydrogen bonds (18 ± 13). The effect of the alternating L and DRI amylin chains is most visible on the stacking of chains which is now stabilized by only (170 ± 16) hydrogen bonds, 151 backbone hydrogen bonds and 19 sidechain hydrogen bonds. connecting L and DRI chains. Hence, as in the previous models, stability of stacking is reduced by about four backbone hydrogen bonds and one sidechain hydrogen bond per L–DRI interface. Note that the number of hydrogen bonds stabilizing the packing of the two protofibrils is with 4 (2) hydrogen bonds stronger than in the (2L-1D-2L)*2 model, leading to a separation of CC-interfaces that with (7.8 ± 0.2 Å) only one Å larger than in the L-amylin fibril and comparable to the pure DRI fibril.

On the other hand, in the second model, 5D*5L, designed by us to probe the effect of alternating L and DRI amylin chains on packing of the two protofibrils, we find a much smaller disturbance of the fibril structure as can be seen from the RMSD and SASA values in Table 1. Here, the two protofibrils preserved most of the interactions seen in their corresponding pure fibril forms, with the number of hydrogen bonds stabilizing the stacking of chains (222 ± 12) comparable to the one found in the L-amylin fibril. However, the number of sidechains stabilizing the packing is again strongly reduced as the loss of Ser29-Ser29 sidechain hydrogen bonds is again only partially compensated by newly formed Ser29–Asn31 hydrogen bonds, which are observed only with a frequency of about 30 % (compared to about 50 % in pure DRI-amylin fibrils). The number of remaining hydrogen bonds (3 ± 2) that stabilize packing of the protofibrils is comparable to that seen in the (2L-1D-2L)*2 model, but the packing is less weakened and the distance between the CC-interfaces with (8.7 ± 1.2 Å) less increased than in the (2L-1D-2L)*2 model. Additional stability comes from *π* − *π* interaction observed between the Phe23 on the DRI protofibril and Tyr37 of the L protofibril. Although these interactions are weaker than hydrogen bonds, they add up and partially compensate for the loss of hydrogen bonds that stabilize in L-amylin fibril the packing of the two protofibrils.

Our third mode, (L/D)*(D/L), is designed to allow us to observe the combined effect of alternating L and DRI-amylin chains on stacking and packing. This arrangement where chain type (L or DRI amylin) changes both between layers and protofibrils leads in terms of RMSD (4.3 ± 0.4) Å), solvent accessible surface area (18370.6 ± 938.2) Å^2^, and distance between the two protofibril (8.4 ± 0.7) Å, to values that lay in between the 5D*5L and the (L/D)*2 model; and shares with both of these models a comparable number of hydrogen bonds (4 ± 6) stabilizing the packing of the two protofibrils. Note that the number of hydrogen bond stabilizing the stacking of chains is with (188 ± 6) much higher than in the (L/D)*2 model (170 ± 17), *i.e.*, the staggering of chains increases the stacking stability but does not affect the packing of protofibrils. This is because unlike in the previous models the *β*2-strands of the two chains in each layer are now parallel instead of anti-parallel, adding not only to stability of the protofibril packing, but by stabilizing the U-shaped geometry of the chains eases also their alignment and stacking.

Hence, in cases where L and DRI strands appears in alternating fashion, *i.e.*, the fibrillar assemblies (L/D)*2 and (L/D)*(D/L), the stacking is reduced by at least four backbone hydrogen bonds and one sidechain hydrogen bond per L–DRI interface, and packing is reduced by four sidechain hydrogen bonds to at most one hydrogen bond connecting two chains on opposite protofibrils (see Table 1). As the latter is a 50 % reduction in number of hydrogen bonds stabilizing the packing of protofibrils, but the former only a (15–20) % reduction in stacking-stabilizing hydrogen bonds, main effect is a separation of double fold assemblies into two single fold fibrils. We conjecture that addition of DRI-amylin discourages elongation of amylin fibrils and causes separation into single-fold fibrils.

## 4 Conclusion

The present study is motivated by the question of whether DRI-amylin could provide an alternative to the often used pramlintide whose utility as a drug for targeting type-II diabetes is limited by its short lifetime. DRI-amylin has the long lifetime associated with peptides built from D-amino acids, but in order to function as a drug it needs to reproduce pramlintide’s ability to inhibit or dis-aggregate amylin aggregates. We have therefore studied by means of molecular dynamics simulations the stability of DRI-amylin fibrils and how insertion of DRI-amylin alters the aggregation of regular L-amylin. We find that fibrils made of DRI amylin are only marginally less stable than the L-amylin fibrils. The difference is due to the loss of about 20 hydrogen bonds (about two per chain), both backbone and sidechain hydrogen bonds. The loss of two backbone hydrogen bonds connecting residue at opposite sides of the *β*-turn region with corresponding residues on a neighbouring layer (His18–Ser19 and Gly24–Ala25) reduces the stability of stacking of chains within each protofibril in DRI-amylin fibrils. Adding to the loss of stacking stability is the halving of the number sidechain hydrogen bonds connecting residues Asn31 on neighbouring layers, as in DRI-amylin fibrils, this residue can now also form sidechain hydrogen bond with Ser29 on the opposite protofibril. These Ser29–Asn31 hydrogen bonds partially compensate for the loss of the Ser29–Ser29 hydrogen bonds that connect the two protofibrils in L-amylin but are not found in DRI-amylin. The net effect is a reduction of packing stabilizing hydrogen bonds by about 25 %, which not so much increases the separation of protofibrils but changes the twist angle of the *β*2-strands that form the interface between the two protofibrils.

Presence of DRI amylin affects the elongation and the stability of L-amylin fibrils by shifting the backbone hydrogen bonds at the L–DRI interface by one residue toward the C-terminus, leading here to a loss of 4 backbone hydrogen bonds in the *β*-turn region and at the C-terminus. Lost are at the L–DRI interface not only the Ser29–Ser29 sidechain hydrogen bonds that stabilize packing of protofibrils in L-amylin but also the Ser29–Asn31 sidechain hydrogen bonds that in DRI-amylin stabilize the packing of protofibrils, partially compensating for the loss of the Ser29-Ser29 hydrogen bonds. This perturbation of hydrogen bonding spreads from the L–DRI interface to successive layers, easing separation of the two protofibrils. Hence, while incorporation of DRI-chains weakens the stacking of chains in each protofibril, the main effect is the reduction of the packing stability between the two protofibrils.

While our simulations indicate that presence of DRI-amylin will inhibit elongation of (L) amylin fibrils and reduce their stability, further studies are needed to explore the kinetics of this process and whether it leads indeed to inhibition of fibril formation. Besides such direct applications, our investigation suggests also that insertion of DRI-proteins in L-assemblies may be an alternative way to mutations to probe the role of such hydrogen bonds in supramolecular assemblies or aggregates. Despite the only marginal structural differences, there are distinct differences in the hydrogen bond pattern in aggregating DRI-peptides, a fact that we used in the present study to point out some key interactions that stabilizes amylin fibril geometries. In particular, Asn ladders (Asn14, Asn22 and Asn31) and Ser–Ser bifurcated hydrogen bonds seems to play an important role in stabilizing L-amylin fibrils. We speculate that the retention of Asn31 in pramlintide contributes to the observed co-aggregation with amylin, and that the effectiveness of the drug could be increased by mutating this residue or replace it with the D-isomer.

## Supporting information

Supplementary Info as refered to in the manuscript

## Acknowledgement

We thank Elliott K. Vanderford for help at early stages of this project. The simulations in this work were done using the SCHOONER cluster of the University of Oklahoma, the Blue Waters sustained - petascale computing project, and XSEDE resources allocated under grant MCB160005 (National Science Foundation). We acknowledge financial support from the National Institutes of Health under grant GM120578.

## Supporting Information Available

Additional tables and figures are provided in supplementary file.

