## Supplementary Info as refered to in the manuscript for "D-Retro Inverso (DRI) Amylin and the Stability of Amylin Fibrils"

### 1 Supplementary Tables

Supplementary Table 1: Twist angle for L-amylin and DRI-amylin.

| | $\beta$1-Pr1 (°) | | $\beta$1-Pr2 (°) | |
|---|---|---|---|---|
| | L-amylin | DRI-amylin | L-amylin | DRI-amylin |
| Run-1 | -12.8 ± 2.9 | 10.7 ± 2.5 | -5.0 ± 3.8 | 15.5 ± 2.4 |
| Run-2 | -23.7 ± 5.4 | 9.3 ± 2.7 | -16.8 ± 2.9 | 12.5 ± 3.4 |
| Run-3 | -16.1 ± 2.6 | 15.7 ± 2.8 | -15.9 ± 2.9 | 13.1 ± 2.6 |
| Avg. | -17.5 (6.0) | 11.9 (3.8) | -12.6 (6.3) | 13.7 (3.1) |
| | $\beta$2-Pr1 (°) | | $\beta$2-Pr2 (°) | |
| | L-amylin | DRI-amylin | L-amylin | DRI-amylin |
| Run-1 | -3.9 ± 1.9 | 7.7 ± 2.0 | -3.2 ± 1.9 | 6.4 ± 2.2 |
| Run-2 | -2.7 ± 2.1 | 4.4 ± 2.0 | -3.2 ± 2.2 | 4.1 ± 1.9 |
| Run-3 | -2.2 ± 2.4 | 9.5 ± 2.0 | -4.5 ± 3.5 | 13.4 ± 2.5 |
| Avg. | -2.9 (2.3) | 7.2 (2.9) | -3.6 (2.7) | 8.0 (4.5) |

Supplementary Table 2: Face to face contact distances of CC interface double layers of L-amylin and DRI-amylin.

| | L-amylin | | | DRI-amylin | | |
|---|---|---|---|---|---|---|
| | Pr1-L2/Pr2-L2 | Pr1-L3/Pr2-L3 | Pr1-L4/Pr2-L4 | Pr1-L2/Pr2-L2 | Pr1-L3/Pr2-L3 | Pr1-L4/Pr2-L4 |
| L27-G33 | 7.5 (0.3) | 7.7 (0.3) | 7.7 (0.4) | 8.8 (1.9) | 8.2 (1.2) | 8.0 (0.6) |
| S29-N31 | 6.5 (0.4) | 6.8 (0.3) | 7.2 (0.3) | 8.3 (1.7) | 7.4 (0.9) | 7.0 (0.5) |
| N31-S29 | 6.6 (0.4) | 6.9 (0.3) | 7.4 (0.3) | 8.2 (1.3) | 7.4 (0.7) | 6.8 (0.3) |
| G33-L27 | 7.9 (0.5) | 8.0 (0.6) | 8.3 (0.7) | 8.9 (0.9) | 8.4 (0.6) | 8.4 (0.7) |

Values are shown after excluding the first and the last layers from each of the protofibril.

Supplementary Table 3: Structural changes in L-amylin (L), DRI-amylin (DRI) and Hybrid Amylin models in terms of average root-mean-square deviation (Å), average Radius of Gyration (Å) and average solvent-accessible-surface-area (Å$^2$).

| | | L | DRI | (4L-1D)*2 | (2L-1D-2L)*2 | (L/D)*2 | 5D*5L | (L/D)*(D/L) |
|---|---|---|---|---|---|---|---|---|
| RMSD | Run-1 | 2.2 ± 0.1 | 3.5 ± 0.2 | 4.1 ± 0.2 | 4.2 ± 0.3 | 6.6 ± 0.3 | 3.3 ± 0.1 | 4.4 ± 0.2 |
| | Run-2 | 4.2 ± 0.5 | 3.3 ± 0.2 | 3.8 ± 0.2 | 6.2 ± 0.2 | 4.1 ± 0.2 | 5.2 ± 0.1 | 4.0 ± 0.5 |
| | Run-3 | 2.6 ± 0.3 | 4.2 ± 0.2 | 3.1 ± 0.2 | 4.7 ± 0.2 | 4.7 ± 0.3 | 2.8 ± 0.4 | 4.4 ± 0.2 |
| | Avg. | 3.0 (0.9) | 3.7 (0.4) | 3.7 (0.5) | 5.0 (0.9) | 5.1 (1.1) | 3.8 (1.1) | 4.3 (0.4) |
| Rg | Run-1 | 22.0 ± 0.2 | 22.4 ± 0.1 | 23.2 ± 0.2 | 22.2 ± 0.2 | 22.0 ± 0.3 | 22.4 ± 0.1 | 22.3 ± 0.3 |
| | Run-2 | 22.8 ± 0.3 | 22.0 ± 0.1 | 22.4 ± 0.2 | 21.8 ± 0.2 | 22.4 ± 0.3 | 22.8 ± 0.2 | 21.6 ± 0.2 |
| | Run-3 | 22.3 ± 0.2 | 22.3 ± 0.2 | 22.6 ± 0.1 | 22.9 ± 0.2 | 21.4± 0.4 | 22.8 ± 0.1 | 22.3 ± 0.2 |
| | Avg. | 22.4 (0.4) | 22.2 (0.2) | 22.7 (0.4) | 22.3 (0.5) | 21.9 (0.5) | 22.7 (0.2) | 22.1 (0.4) |
| SASA | Run-1 | 16469 ± 347 | 18180 ± 520 | 18389 ± 236 | 17594 ± 503 | 20285 ± 613 | 17176 ± 353 | 19158 ± 604 |
| | Run-2 | 18283 ± 515 | 17257 ± 383 | 17551 ± 382 | 18197 ± 477 | 19019 ± 538 | 18263 ± 373 | 17410 ± 595 |
| | Run-3 | 17413 ± 407 | 17724 ± 437 | 17132 ± 410 | 18110 ± 404 | 18213 ± 797 | 16844 ± 313 | 18544 ± 590 |
| | Avg. | 17388 (856) | 17720 (587) | 17691 (629) | 17967 (534) | 19172 (1077) | 17428 (699) | 18371 (938) |

RMSD values are with respect to the start configuration and calculated over the last 80 $ns$. Only backbone atoms are considered, and the first seven residues of the flexible N-terminus not taken into account.

#### 2 Supplementary Figures

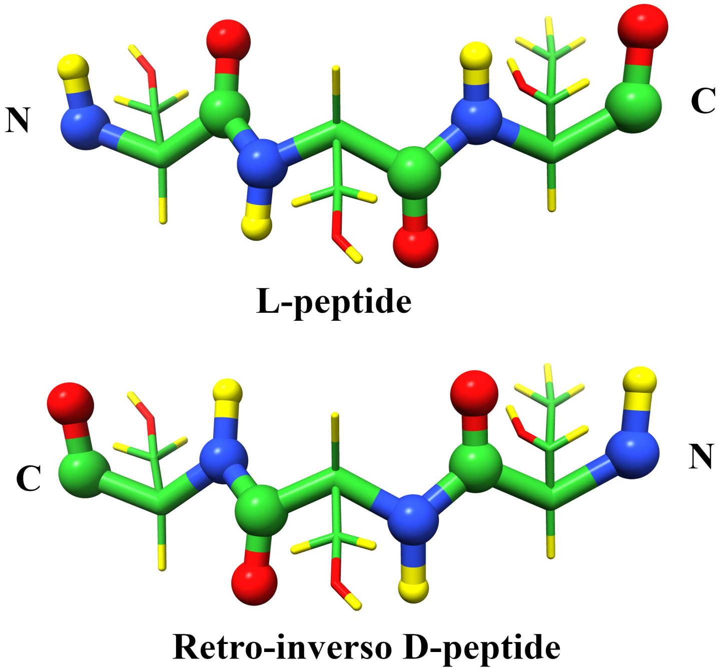


Supplementary Figure 1: Backbone switching to convert L-peptide into its retro-inverso DRI form made of D-amino acids. The backbone atoms are shown as spheres using the color code: N-blue, HN-yellow, C-green and O-red.

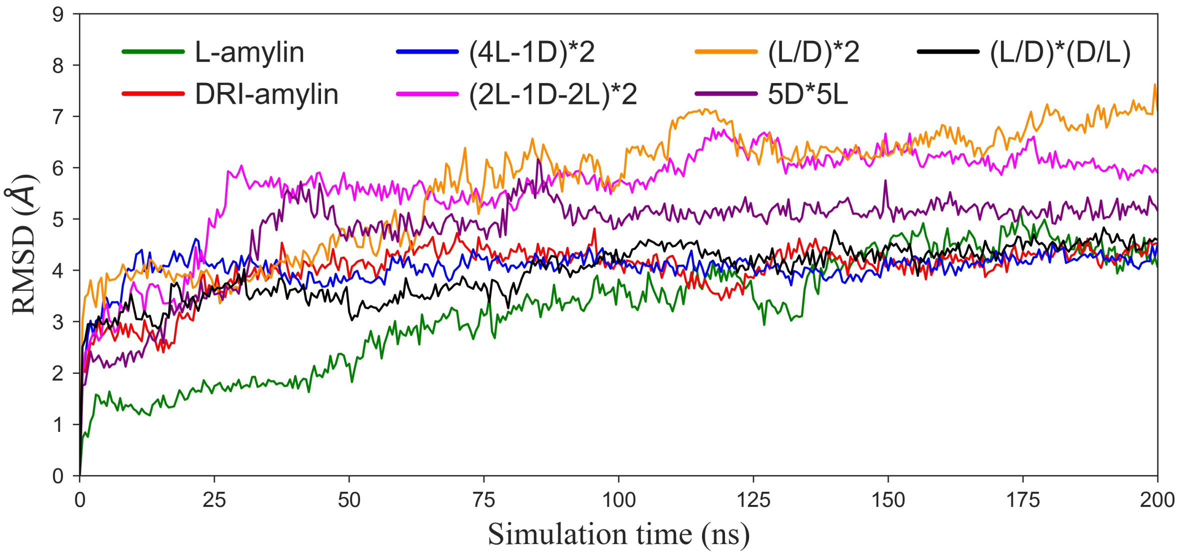


Supplementary Figure 2: Time evolution of root-mean-square deviation (RMSD) of L-amylin, DRI-amylin and L & DRI mixtures. RMSD values are calculated with respect to the starting configuration considering only backbone atoms. Due to higher fluctuation of first 7 residues, they were ignored while calculation of RMSD. Time evolution of RMSD is only shown for the trajectories with the highest fluctuations for the various amylin assemblies.

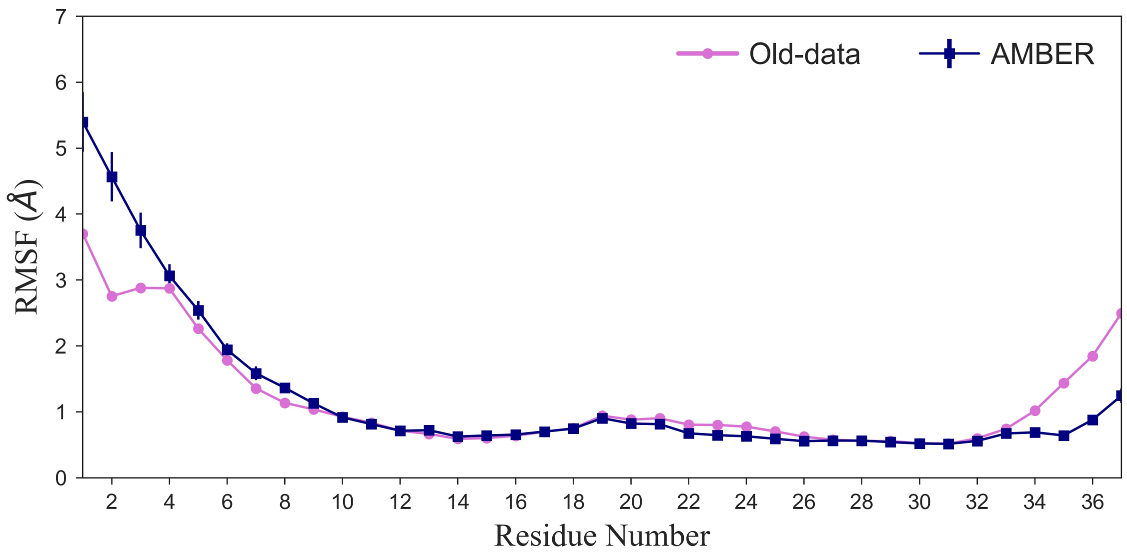


Supplementary Figure 3: Comparison of Root-mean-square fluctuation (RMSF) of the C$\alpha$ atoms for AMBER force-field with Berhanu *et al.* (2014). Values are averaged over all monomers and the three trajectories.

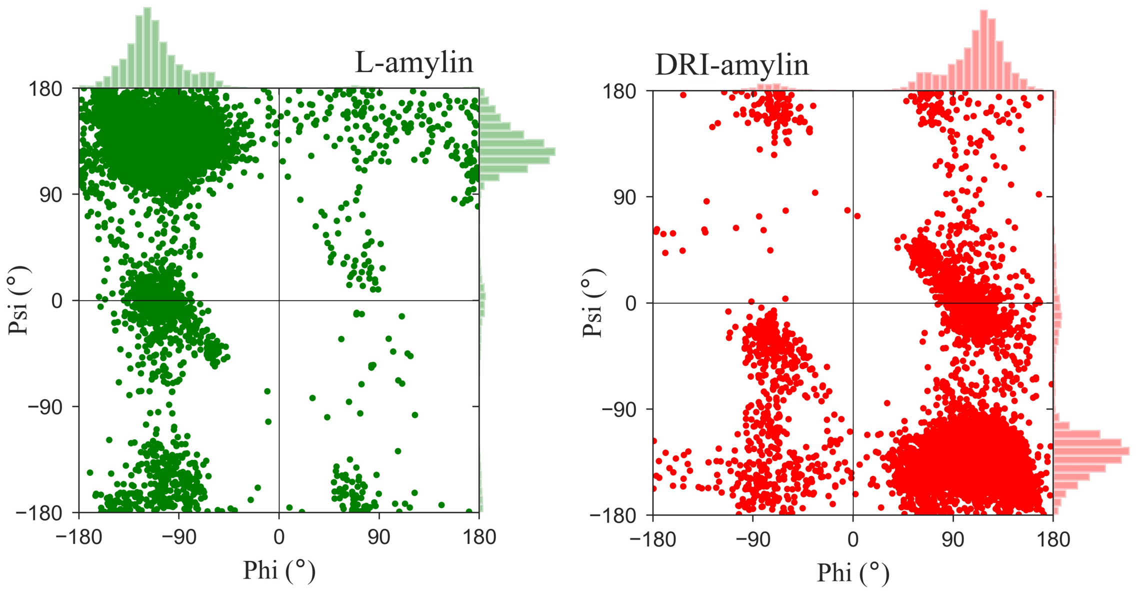


Supplementary Figure 4: Ramachandran plot for L-amylin and DRI-amylin.

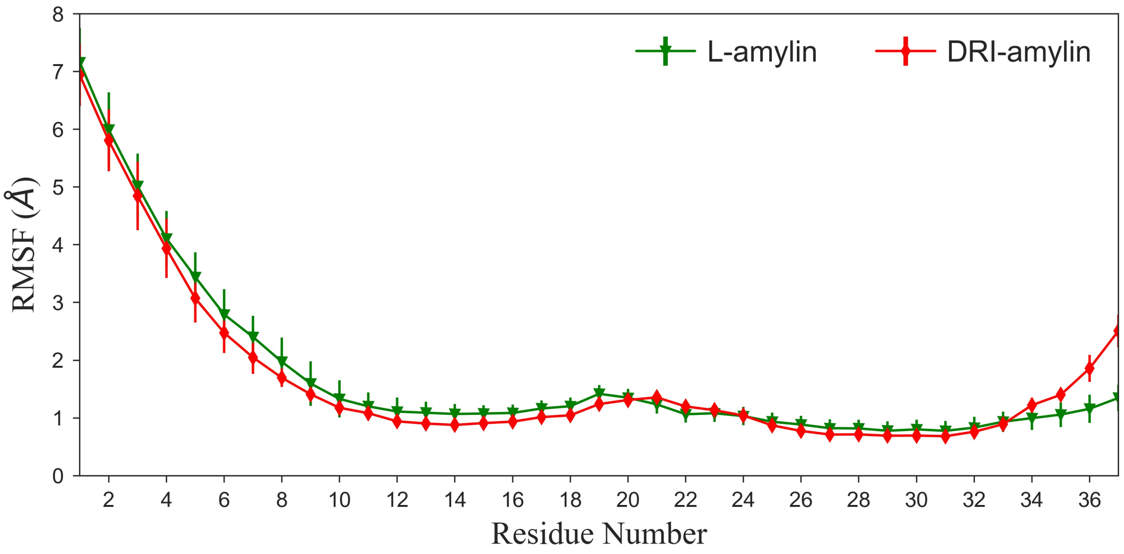


Supplementary Figure 5: Root-mean-square fluctuation (RMSF) of the C$\alpha$ atoms for L-amylin and DRI-amylin. Values are averaged over all monomers and the three trajectories.
